# Increases in serum corticosterone level and diuresis induced by the selective kappa opioid receptor (KOR) agonist U50,488H are unaffected by KOR phosphorylation

**DOI:** 10.64898/2026.07.10.737784

**Authors:** Pingwei Zhao, Kathryn Bland, Sanjana Khandeshi, Peng Huang, Lee-Yuan Liu-Chen

**Author notes:** **Correspondence should be addressed to** Lee-Yuan Liu-Chen, Ph.D., Center for Substance Abuse Research, Temple University Lewis Katz School of Medicine, 3500 North Broad Street, MERB 851, Philadelphia, PA 19140, USA. **Availability of data and materials** Data will be available from the corresponding author upon reasonable request.

## Abstract

**Purpose:** We previously showed that mice expressing a phosphorylation-deficient kappa opioid receptor mutant (K4A) exhibited reduced U50,488H-induced anti-scratching tolerance in males and reduced conditioned place aversion in females, without changes in acute anti-scratching or hypo-locomotor effects. Here, we examined whether K4A mutations, which markedly diminish β-arrestin-mediated signaling, alter U50,488H-induced increases in serum corticosterone and urine output.

**Methods:** K4A and wildtype mice received U50,488H (5 mg/kg, s.c.) or saline. Serum corticosterone was measured by ELISA 1 h later. Urine was collected for 1 h beginning 10 min after injection.

**Results:** U50,488H increased serum corticosterone to similar levels in wildtype and K4A mice of both sexes. Basal corticosterone levels were higher in females than males regardless of genotype. U50,488H also significantly increased urine output in both sexes, with no genotype differences. However, the increase in urine output was greater in males than females.

**Conclusions:** KOR phosphorylation and associated β-arrestin-mediated signaling are not required for U50,488H-induced increases in serum corticosterone or diuresis in either sex. These findings also demonstrate, for the first time, that KOR activation produces greater diuresis in male than female mice.

## INTRODUCTION

Activation of kappa opioid receptor (KOR), coupled to G_i/o_ proteins, produces analgesia, anti-pruritic effect, and water diuresis (1-4). KOR agonists may be useful for prevention and treatment of substance use disorders [for a review, (5)]. However, KOR agonists also cause sedation, dysphoria, psychotomimesis, and increased corticosteroid levels (1, 2, 6-8), limiting their clinical development.

Activation of G protein-coupled receptors (GPCRs) initiates G proteins-mediated signaling, receptor phosphorylation by GPCR kinases, and recruitment of arrestins, leading to receptor regulation and arrestins-mediated signaling. G proteins and arrestins initiate distinct signaling pathways and produce different in vivo responses. Biased agonism has attracted considerable interest, although its validity remains debated [for example, (9-12)]. This concept suggests that G protein- or arrestins-biased agonists may produce therapeutic effects with reduced adverse effects. For KOR, analgesic and anti-scratching effects are generally attributed to G protein signaling, whereas the roles of G protein versus β-arrestin signaling in dysphoria, hypolocomotion, and motor impairment remain controversial [for a review, (13)].

We previously identified U50,488H-induced KOR phosphorylation sites (S356, T357, T363, and S369 in the C-terminal domain) and generated a knock-in mouse line expressing a phosphorylation-deficient KOR mutant (K4A; S356A/T357A/T363A/S369A) (14, 15). K4A mutations abolished U50,488H-induced KOR phosphorylation. Thus, the K4A mutations essentially eliminate arrestins-mediated signaling. K4A mutations attenuated U50,488H-induced tolerance in anti-scratch effect in males only, and reduced U50,488H-promoted conditioned place aversion (CPA) in females only (15). In contrast, K4A mutations did not affect acute U50,488H-induced anti-pruritic and hypo-locomotor effects (15). Whether KOR phosphorylation and related arrestin signaling contribute to U50,488H-induced corticosterone elevation or diuresis remains unknown.

Here, we compared the effects of U50,488H in wildtype and K4A male and female mice on serum corticosteroid and diuresis. A dose of 5 mg/kg (s.c.) was selected because it produces maximal anti-scratch and formalin analgesic effects in mice for the following reasons. In mice, this dose produces maximal analgesic effects in the second phase of the formalin test and also maximal anti-scratch effects against compound 48/80-induced scratch (16, 17). In rats, 3 and 10 mg/kg U50,488 (s.c.) enhanced diuresis to similar extents (1, 18, 19), and U50,488H elevated plasma corticosteroid levels and at 2.3 mg/kg caused a 3x increase (1). U50,488H showed similar potencies in wildtype and K4A mice in the anti-scratching test (15) .

## MATERIALS AND METHODS

### Materials

U50,488H was obtained from the NIDA Drug Supply Program (Research Triangle Institute, NC). Filter papers (grade no. 1) were purchased from Advantec MFS (Dublin, CA). A corticosterone ELISA kit (Cat# KGE009) was purchased from R&D Systems (Minneapolis, MN).

### Animals

Adult male and female wildtype C57BL/6N mice and homozygous K4A mice on a C57BL/6N background (25–40 g, 5–8 months old) were used. Wildtype mice were purchased from Jackson Laboratory (Bar Harbor, ME). The K4A knockin line was customed generated by homologous recombination (Cyagen, Santa Clara, CA), bred in-house, and genotyped as we described previously (15). K4A mice were cryopreserved at Jackson Laboratory [#415239, Oprk1tm2.2(K4A)Llch]. Mice were housed under standard conditions (22°C, 50–60% humidity, 12:12 h light/dark cycle) with food and water ad libitum. All procedures were approved by the Temple University Institutional Animal Care and Use Committee.

### Blood collection and corticosterone determination

Mice were habituated to handling and injection once a day for 3 days. Mice then received U50,488H (5 mg/kg, s.c.) or saline (both 10 μL/g body weight). One hour later, 50–100 μL blood was collected from the submandibular vein according to NIH guidelines for survival blood collection (http://oacu.od.nih.gov/ARAC/survival.pdf). One week later, mice received the alternate treatment in a counterbalanced design. Serum corticosterone was measured using a commercial ELISA kit following published procedures (20, 21). Serum samples were processed according to the manufacturer’s instructions, and absorbance was measured at 450 nm with background subtraction at 550 nm. Concentrations were calculated from standard curves using Prism 10 (GraphPad Software).

### Urine output determination

Following ≥1 h habituation, mice received U50,488H (5 mg/kg, s.c.) or saline and were returned to home cages for 10 min. Mice were then placed on pre-weighed filter papers in plexiglass cylinders for 1 h. Urine output was determined by filter paper weight gain after removal of feces and normalized to body weight. Treatments were separated by a 72-h washout period.

### Data analysis

Data are presented as mean ± S.E.M. and were analyzed using two-way ANOVA.

## RESULTS

### Effects of U50,488H on serum corticosterone levels

Two-way ANOVA revealed a significant main effect of treatment, but no effect of genotype or treatment × genotype interaction. Thus, U50,488H (5 mg/kg, s.c.) significantly increased corticosterone levels in male and female wildtype and K4A mice, with no genotype differences (Fig. 1A,B).

**Figure 1.**
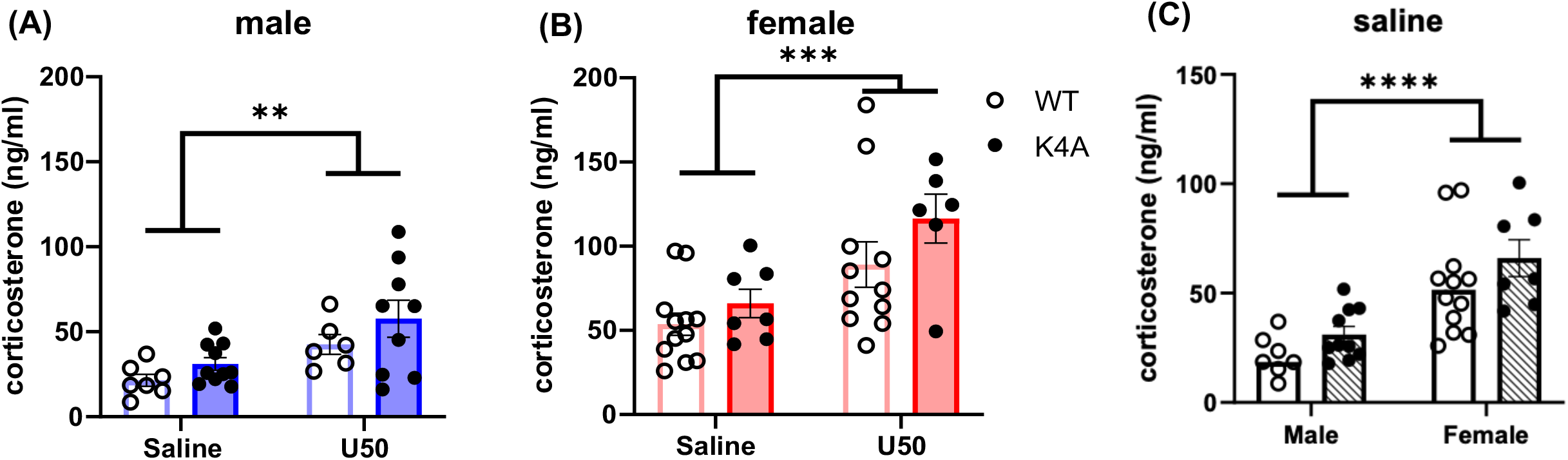
U50,488H increased serum corticosterone levels in (A) male and (B) female wildtype and K4A mice and there is no genotype difference. Male and female wildtype and K4A mice were injected with saline or U50,488H (5 mg/kg, s.c) and blood collected 60 min later. Corticosterone levels in serum were determined with an ELISA kit. Results of 2-way ANOVA analysis showed that there was a significant main effect of treatment in either sex [males, F (1, 28) = 10.80, P=0.0027] [females, F(1, 32) = 13.89, P=0.0007], but no significant main effect of genotype [males, F (1, 28) = 2.923, P=0.0984] [females, F (1, 32) = 2.964, P=0.0948] or genotype x treatment interaction [males, F (1, 28) = 0.1459, P=0.7053] [females, F (1, 32) = 0.4227, P=0.5203]. Data are expressed as mean ± S.E.M. (N = 6–12 per group). **P < 0.01, ***P< 0.001, compared with saline **(C) Basal serum corticosterone levels were higher in female wildtype and K4A mice than male mice**. Data of the saline-treated groups from (A) and (B) were analyzed using 2-way ANOVA, which revealed significant main effects of sex [F (1, 33) = 29.96, P<0.0001] but not genotype [F (1, 33) = 3.247, P=0.0807]. Thus, female mice had higher basal corticosterone levels than male mice of the same genotype. Data are expressed as mean ± S.E.M. (N = 7–12 per group). ****P < 0.0001, compared with male counterparts.

Basal corticosterone levels following saline treatment were compared between sexes (Fig. 1C). Two-way ANOVA showed a significant main effect of sex, but no effect of genotype or sex × genotype interaction. Therefore, female mice exhibited higher corticosterone levels than males. Analysis of U50,488H-induced net increases also revealed no significant effects of genotype or sex.

### Effects of U50,488H on urine output

Two-way ANOVA revealed a significant main effect of treatment, but no effect of genotype or treatment × genotype interaction. Accordingly, U50,488H (5 mg/kg, s.c.) significantly increased urine output in both sexes regardless of genotype (Fig. 2A,B). Net increases in urine output were calculated by subtracting the mean saline value from each U50,488H-treated data point. Two-way ANOVA showed a significant main effect of sex, but no effect of genotype or sex × genotype interaction (Fig. 2C). Thus, U50,488H increased urine output to a greater extent in males than females, independent of genotype.

**Figure 2.**
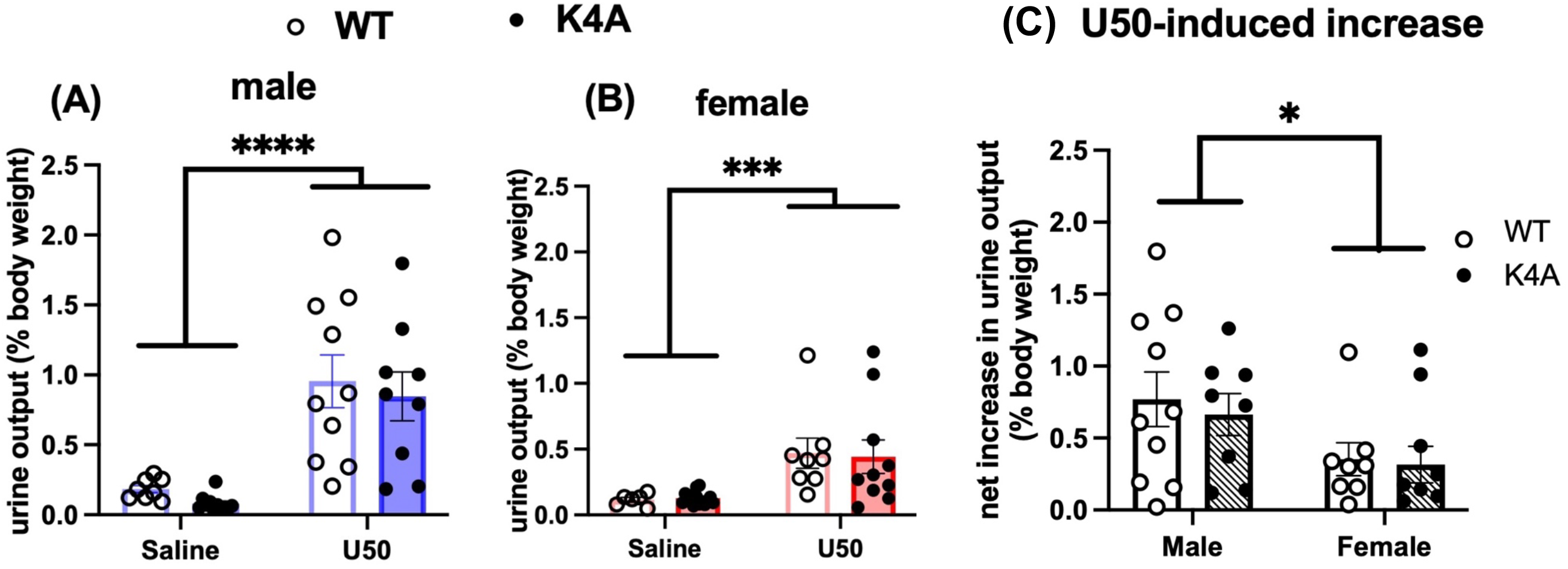
In (A) male or (B) female mice U50,488H increased urine output in WT and K4A mice significantly with no genotype differences. Mice were injected with saline or U50,488H (5 mg/kg, s.c.) and measurement of urine output started 10 minutes after injection for 1 hour. Data for males and females were analyzed separately with 2-way ANOVA, which showed significant main effects of treatment [males, F (1, 34) = 34.16, P<0.0001] [females, F (1, 34) = 14.71, P=0.0005] but no significant main effects of genotype [males, F (1, 34) = 0.6443, P=0.4277] [females, F (1, 34) = 0.006825, P=0.9346] or treatment x genotype interaction [males, F (1, 34) = 0.0002200, P=0.9883] [females, F (1, 34) = 0.04719, P=0.8293]. Data are expressed as mean ± S.E.M. (N = 6-10 per group). ***P<0.001, ****P<0.0001 compared with saline of the same group. **(C) U50,488H-induced net increase in urine output is greater in males than in females**. Net increases were calculated by subtracting the mean of each saline group from individual values of the same sex-genotype group. Data were analyzed with 2-way ANOVA, which shows a significant main effect of sex [F (1, 32) = 6.276, P=0.0175] but not genotype [F (1, 32) = 0.2262, P=0.6376] or sex x genotype interaction [F (1, 32) = 0.05151, P=0.8219]. Data are expressed as mean ± S.E.M. (N= 8-10/group). *P<0.05 between males and females.

## DISCUSSION

This is the first study to examine the role of KOR phosphorylation in KOR agonist-induced increases in serum corticosterone and diuresis. U50,488H increased serum corticosterone levels and urine outputs similarly in wildtype and K4A mice of both sexes. Females exhibited higher basal corticosterone levels, whereas males showed greater U50,488H-induced net increases in urine output independent of genotype. These findings indicate that KOR phosphorylation and the subsequent β-arrestin-mediated signaling are not involved in U50,488H-induced increases in serum corticosterone or diuresis, which are likely mediated by G protein signaling.

### Signaling underlying KOR agonist-induced diuresis

Our findings are consistent with those of Inan, Lee (18) who showed that in the dose ranges producing effective anti-scratch effects U50,488H (1-10 mg/kg) and nalfurafine (5-20 μg/kg) increased urine output dose-dependently in rats. We have shown that *in vivo* U50,488H (5 mg/kg), but not nalfurafine (20 μg/kg), promotes KOR phosphorylation in mouse brains and KOR internalization in the mouse ventral tegmental area (22). As both compounds produced effective diuresis, KOR phosphorylation and internalization, proxies of β-arrestin-mediated signaling, are unlikely to be required. In contrast, triazole 1.1 and triazole 187, demonstrated to be G protein-biased *in vitro*, produced diuresis, albeit with lower maximal effects than U50,488H (23), suggesting possible β-arrestin involvement. Differences in experimental approaches may account for this discrepancy.

### Signaling underlying KOR-mediated *in vivo* effects

Previous studies have demonstrated that KOR-mediated analgesic and anti-pruritic effects are mediated by G protein signaling [for reviews, see (13, 24, 25)], based on findings from mutant mouse lines (GRK3-/-, β-arrestin2-/-, p38α MAPK-/-, K4A), distinct KOR agonists, and G protein-biased agonists. Our data indicate that KOR-induced increases in serum corticosterone and urine output are also likely mediated by G proteins. In contrast, mechanisms underlying KOR-mediated side effects, especially CPA, remain unresolved. Studies using genetic and pharmacological approaches indicate distinct mechanisms for CPA, hypolocomotion, and motor incoordination. CPA involves GRK3-, p38α MAPK, mTOR, CB1 receptors, and PKC activation, but not β- arrestin2. KOR phosphorylation is required for U50,488H-induced CPA in female but not male mice. Several G protein-biased KOR agonists produce antinociceptive and anti-scratch effects with reduced side effects, but non-identical side effect profile, and some still induce CPA. PKC inhibition impairs novelty-induced hyperlocomotion and reduces motor incoordination, whereas mTOR inhibition, β-arrestin2 deletion, or KOR phosphorylation deficiency does not. Motor incoordination is reduced by PKC inhibition or β-arrestin2 deletion, but unchanged by mTOR inhibition.

### Choice of U50,488H treatment intervals

For corticosterone measurements, blood samples were collected 1 h after U50,488H administration and urine output was measured from 10–70 min post-injection, based on prior studies of corticosterone, diuresis, and KOR phosphorylation. In rats, blood samples for corticosterone measurements were obtained 1 h after U50,488H administration (1, 18). U50,488H-induced KOR phosphorylation peaks at 30–60 min (22).

### U50,488H increases serum corticosterone level

Our finding that U50,488H increases serum corticosterone in mice is consistent with reports in mice, rats, and humans (1, 26-28) Most previous studies were conducted in males; here we extend these findings to females. Basal corticosterone levels were higher in females, consistent with previous reports (29, 30). This sex difference likely reflects reduced negative regulation of the hypothalamic-pituitary-adrenal axis (29, 30) and higher corticosteroid-binding globulin concentrations in females (31).

### U50,488H increases urine output: sex differences

U50,488H and other KOR agonists increase urine output in rats, dogs, monkeys, and humans, and these effects are blocked by KOR antagonists (1, 26, 32-34)[reviewed in (35)]. We extend these findings to mice and females, showing greater U50,488H-induced diuresis in males when normalized to body weight. In contrast, Craft, Ulibarri (19) reported similar responses in male and female rats. The basis for this discrepancy is unclear. KOR agonists have been explored as water diuretics for water retention associated with edema, elevated intracranial pressure, and liver cirrhosis (36-39). Unlike traditional diuretics, they cause little or no loss of K^+^ and Na^+^ in urine. Potential sex differences in humans warrant further investigation.

## Author Contribution Statement

PZ designed research, performed experiments, analyzed data, revised the manuscript.

KB performed experiments and analyzed data.

SK performed experiments.

PH designed research, performed experiments, analyzed data. LYLC conceived of study, analyzed data, wrote the paper.

## Acknowledgments

We thank Dr. Saadet Inan for helpful inputs and Mr. Chongguang Chen for technical assistance.

